# Stress, heritability, tissue type and human methylome variation in mother-newborn dyads

**DOI:** 10.1101/002303

**Authors:** David A. Hughes, Nicole C. Rodney, Connie J. Mulligan

## Abstract

DNA methylation variation has been implicated as a factor that influences inter-individual and inter-tissue phenotypic variation in numerous organisms and under various conditions. Here, using a unique collection of three tissues, derived from 24 mother-newborn dyads from war-torn Democratic Republic of Congo, we estimate how stress, heritability, tissue type and genomic/regulatory context influence genome-wide DNA methylation. We also evaluate if stress-associated variation may mediate an observed phenotype - newborn birthweight. On average, a minimal influence of stress and heritability are observed, while in contrast extensive among tissues and context dependency is evident. However, a notable overlap in heritable and stress-associated variation is observed and that variation is commonly correlated with birthweight variation. Finally, we observe that variation outside of promoter regions, particularly in enhancers, is far more dynamic across tissues and across conditions than in promoters, suggesting that variation outside of promoters may play a larger role in expression variation than variation found within promoter regions.

## Introduction

The methylation of cytosines is an important source of epigenetic variation that varies among tissues, individuals and species (Pai et al. 2011). Cytosine methylation has been demonstrated to correlate with gene expression variation (Pai et al. 2011; Eckhardt et al. 2006) and is thought to play a regulatory function that influences cellular and organismal phenotypes (Waterland and Jirtle 2003). An initial focus on CpG island and promoter methylation has emphasized the role of methylation in influencing expression variation by affecting transcription initiation. However, recent genome-wide methylation profiling has implicated methylation variation in other genomic contexts (Ziller et al. 2013) which may influence phenotypes by affecting transcription elongation, chromosomal stability or chromatin accessibility (Jones 2012).

Additionally, cytosine methylation has exhibited correlated responses to environmental stimuli. Investigations ranging from insects to humans have illustrated epigenetic responses to environmental stimuli that include stress, age, diet, and toxins (Faulk and Dolinoy 2011; Grönniger et al. 2010; Carone et al. 2010). Other studies (Heijmans et al. 2008; Borghol et al. 2012; Mulligan et al. 2012; Radtke et al.) have indicated that the neonatal or periconceptual environment is a crucial and sensitive period of epigenetic programming (Faulk and Dolinoy 2011). These observations are consistent with the “developmental origins of health and disease” hypothesis (DOHaD) (Barker 1990; Gluckman et al. 2010; 2007), arguing that the fetus is most sensitive to external stimuli during cellular differentiation, growth and development, i.e. the time points during which DNA methylation profiles are being established. As a result, the epigenetic response to negative stimuli could prove maladaptive and lead to disease later in life. For example, newborn methylation variation at the glucocorticoid receptor *NR3C1* has been previously associated with maternal stress and newborn birthweight (Mulligan et al. 2012) while in other studies *NR3C1* variation correlates with other stressors and phenotypes such as child abuse, suicide, depression, birthweight and intimate partner violence (Oberlander et al. 2008; McGowan et al. 2009; Mulligan et al. 2012; Radtke et al.; Filiberto et al. 2011).

The heritability of methylation variation remains a topic of debate. It is apparent that a small proportion of methylated cytosines are influenced by nearby *cis* acting genetic variants (Gertz et al. 2011; Fraser et al. 2012; Bell et al. 2011; Gibbs et al. 2010; Zhang et al. 2010). The nature and significance of what has been called methylation ‘soft-inheritance’ (Richards 2006) or transgenerational epigenetic inheritance (Rakyan and Whitelaw 2003; Jirtle and Skinner 2007; Morgan and Whitelaw 2008) remains to be determined (Grossniklaus et al. 2013; Richards 2006). Transgenerational epigenetic inheritance (TEI) refers to non-genetic inherited effects. These are methylation modifications that correlate or correspond to environmental conditions that are subsequently passed to future generations. TEI has been observed in numerous studies and under numerous environmental conditions including diet, pollution and stress (Cortessis et al. 2012; Jirtle and Skinner 2007; Feil and Fraga 2012; Terry et al. 2011). A well-studied example comes from observations on the influence of diet on coat color variation in mice (Waterland and Jirtle 2003; Dolinoy et al. 2006). However, it has been argued that to be a genuine TEI variant, such variation must be maintained through the F_3_ generation because the F_0_ mother, the F_1_ offspring and the F_2_ germ line have all experienced the same or similar environment(s). Currently, there are few studies of TEI consistent with this definition (Anway et al. 2005). Due to a unique set of samples tissues, we are well positioned to evaluate the role that environment has on methylation inheritance between F_0_ mothers and F_1_ newborns.

In this study, we evaluate the influences of environmental stimuli (maternal stress), heritability, tissue type and genomic context on genome wide methylation variation in three tissues - maternal venous blood, cord blood and placenta. Our samples are of maternal and fetal origin, collected from women in Goma, Democratic of Congo (DRC) - a region of ongoing war and extreme physical and psycho-social stress to women. These women have experienced a range of physical and emotional stressors including rape, physical abuse, and refugee status. In addition to the tissue samples, detailed ethnographic information was collected through open-ended interviews with each mother in order to develop emic, i.e. culturally relevant, measures of stress (Mulligan et al. 2012). We investigate the associations of maternal stress with maternal and newborn methylome variation and then query their potential influence on newborn birthweight.

## Results

### Maternal Stress

Maternal stress was quantified by building a simple additive model (as in Mulligan et al. (Mulligan et al. 2012)) using 13 war-related stress questions (Table 1). In addition to the data on war stressors, 11 medical and educational variables were also acquired, including maternal age, maternal weight and household education level (the sum of husband and wife education level) (Table 1). We evaluated the potential dependences of age, weight and education on maternal stress by correlation and regression analyses. Correlations between age and stress (r = −0.482, p = 0.0202) as well as education and stress (r = −0.734, p = 8.785e-05) were observed. In a two factor ANOVA the proportion of explained variation (η^2^, see Materials and Methods) for age and education were 23.1% (F-test p-value = 3.31e-3) and 35.2% (F-test p-value = 5.4e-4), respectively. A model including interaction between age and education does not improve the fit of the data. In short, these results indicate that, in this population, both the age of the woman and the education level of her household influence maternal stress.

**Table 1.** Questions Included in Trauma Surveys

| Stressors | Other Factors |
| --- | --- |
| Have you been a refugee in the past? | Age at first pregnancy? |
| Are you currently a refugee? | Maternal age |
| Has a family member been killed in a war? | Maternal weight |
| Have you been raped in the past? * | Parity |
| Did you have a child as a result of rape in the past? * | Labor duration |
| Is your current pregnancy the result of rape? * | Do you drink? |
| Were you raped during your current pregnancy? * | Gestational age |
| Have you ever been kidnapped? | Birthweight |
| Has a parent been raped? | Sex |
| Are you a child of rape? | Mother's education level |
| Do you currently reside in a conflict area? | Father's education level |
| Have you been physically abused? |  |
| Have you been emotionally abused? |  |

A simple additive model of stress was created by simply adding 1, to an individual's stress level each time they would say "yes" to one of the 13 Stressors questions. Our model of rape only includes the four stressors containing a \*. Other Factors includes additional information that was collected and evaluated.

### Stress influences birthweight

Many investigations with varying types of stressors have consistently observed a correlation between maternal stress and newborn birthweight (Eskenazi et al. 2007; Rondó et al. 2003; Khashan et al. 2008; Zhu et al. 2010; Orr et al. 1996; Collins et al. 2004; Lee et al. 2011; Feeley et al. 2011) and a recent World Health Organization report concluded that, globally, women who have been physically or sexually abused are 16% more likely to have an offspring of low birth weight (WHO et al. 2013). In our study, we observed a significant correlation between maternal stress and newborn birthweight (Fig. 1A, r = −0.627, p = 0.001), with stress explaining an estimated (η^2^) 39.3% of the total variation in birthweight. In an effort to further simplify our model of stress and determine if the timing of a stressful event influences birthweight, we used the relative time since the mother’s last personal experience of rape and observed a similar effect (p = 0.0383), explaining 33.7% of the birthweight phenotype (Fig. 1B). Additionally, household education level (p = 0.0005, Fig. 1C) and maternal age (p = 0.0345, Fig. 1D) also correlated with birthweight. Jointly, the inter-correlations of stress, age, and education with birthweight confound the identification of a single explanatory variable. However, we hypothesize that a woman’s age and household education influence her level of stress and it is the physical properties of stress that are influencing birthweight - as observed in the references sited above. Similar correlations were observed when modeling these factors’ effects on gestational age, but we chose to focus on birthweight given the robust and replicable association between this measure and later disease(Barker et al. 2002; Gluckman et al. 2008; Innes et al. 1999) and because all gestational ages were determined ad hoc at the time of delivery.

**Figure 1.**
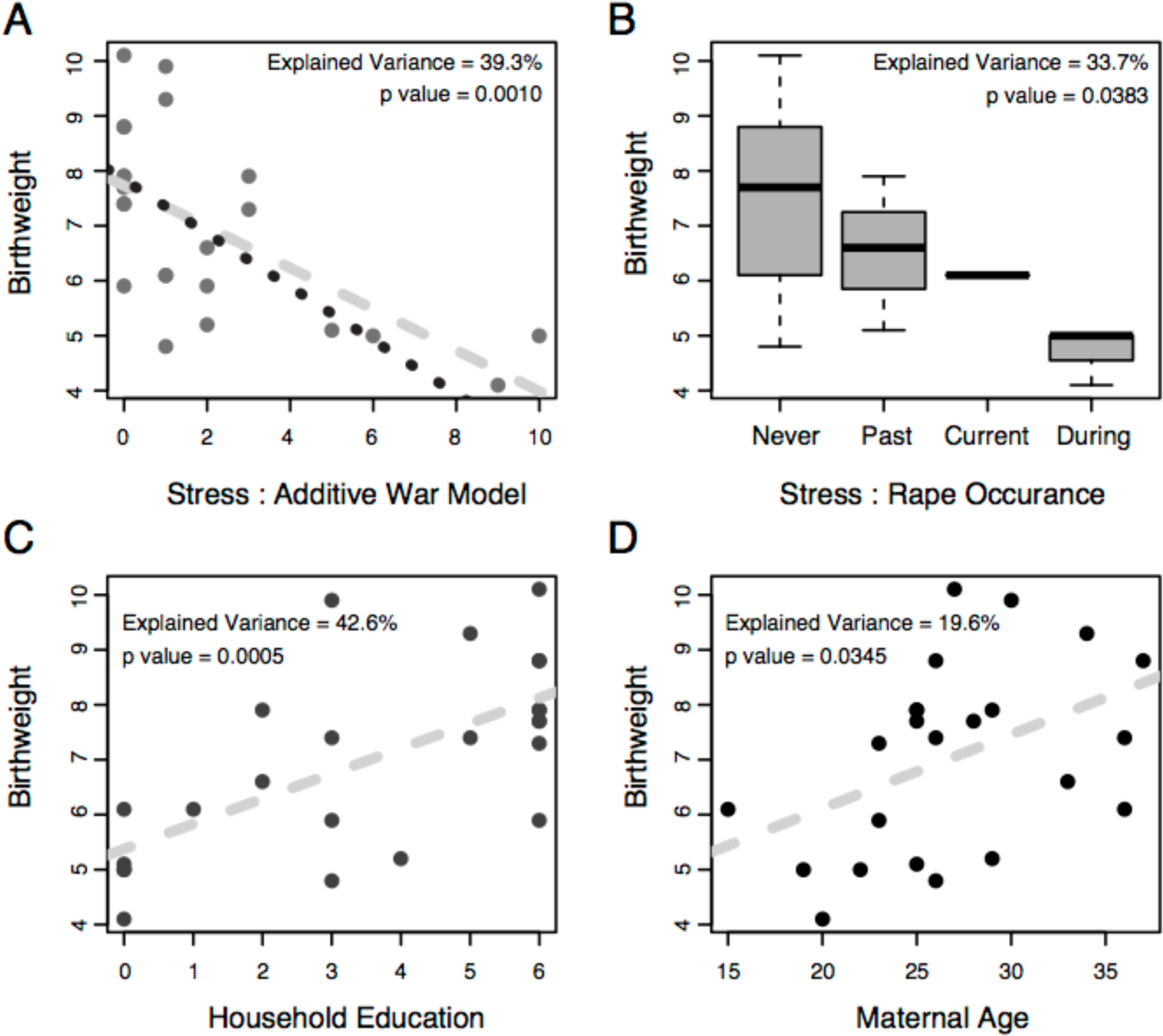
Correlation analysis for newborn birthweight and (A) war stress, (B) rape occurrence, (C) household education and (D) maternal age. The dashed line in plots (A), (C) and (D) represents the linear regression for all the data points. The dotted line in plot (A) is the linear regression after removing the two most extreme samples. Plot (B) is a box plot of birthweights after partitioning women into their most recent experience of rape. ‘Never’ represents women who were never raped, ‘Past’ represents women who have had an experience of rape at some time point in the past. ‘Current’ represents women whose current pregnancy is the result of rape. ‘During’ represents women who were raped during their current pregnancy. The proportion of variance explained (**η^2^**) for each explanatory variable and its F-test p-value are denoted in each plot.

### Methylation data

Genome wide methylation variation was interrogated at 485,577 CpG sites across the genome using the Illumina HumanMethylation450 BeadChip technology. Ninety-one percent of all probes were significantly detected above background signal in all 72 (24 dyads*3 tissues) samples. An additional 10,423 probes located on the sex chromosome were removed from the analysis, yielding 431,048 high quality autosomal probes for analysis. The proportion of methylation at each CpG was summarized with the β statistic, defined as the ratio of methylated probe intensities over the sum of intensity values at both methylated and un-methylated probes for that site.

### Global methylation variation

Global methylation was summarized into a single statistic for each individual by simply averaging methylation (beta values) across all CpG sites - separated by tissue. A significant difference in global methylation between tissues was observed (Fig. 2A). The placenta is hypomethylated (mean beta = 0.451) relative to both venous blood (mean beta = 0.507, Welch two sample t-test p-value = 2.2e-16) and cord blood (mean beta = 0.502, p-value = 2.2e-16) (Fig. 2A). Cord blood and venous blood tissues are only modestly different from each other but significantly so (p-value = 0.0156). Additionally, there is a visible decrease in among-individual variation in venous blood samples relative to the other tissues (Fig. 2A). We quantified among individual variation by calculating the coefficient of variation or Eta (η), where Eta is defined as the standard deviation among individuals divided by the mean. Venous tissue has a η of 0.007, 2.6 times smaller than that observed in either of the other tissues (0.018) suggesting a potential difference in the mechanistic regulation of methylation variation among the tissues. However, this observation could reflect greater levels of cell type heterogeneity across the placenta and cord blood samples, relative to venous blood.

**Figure 2.**
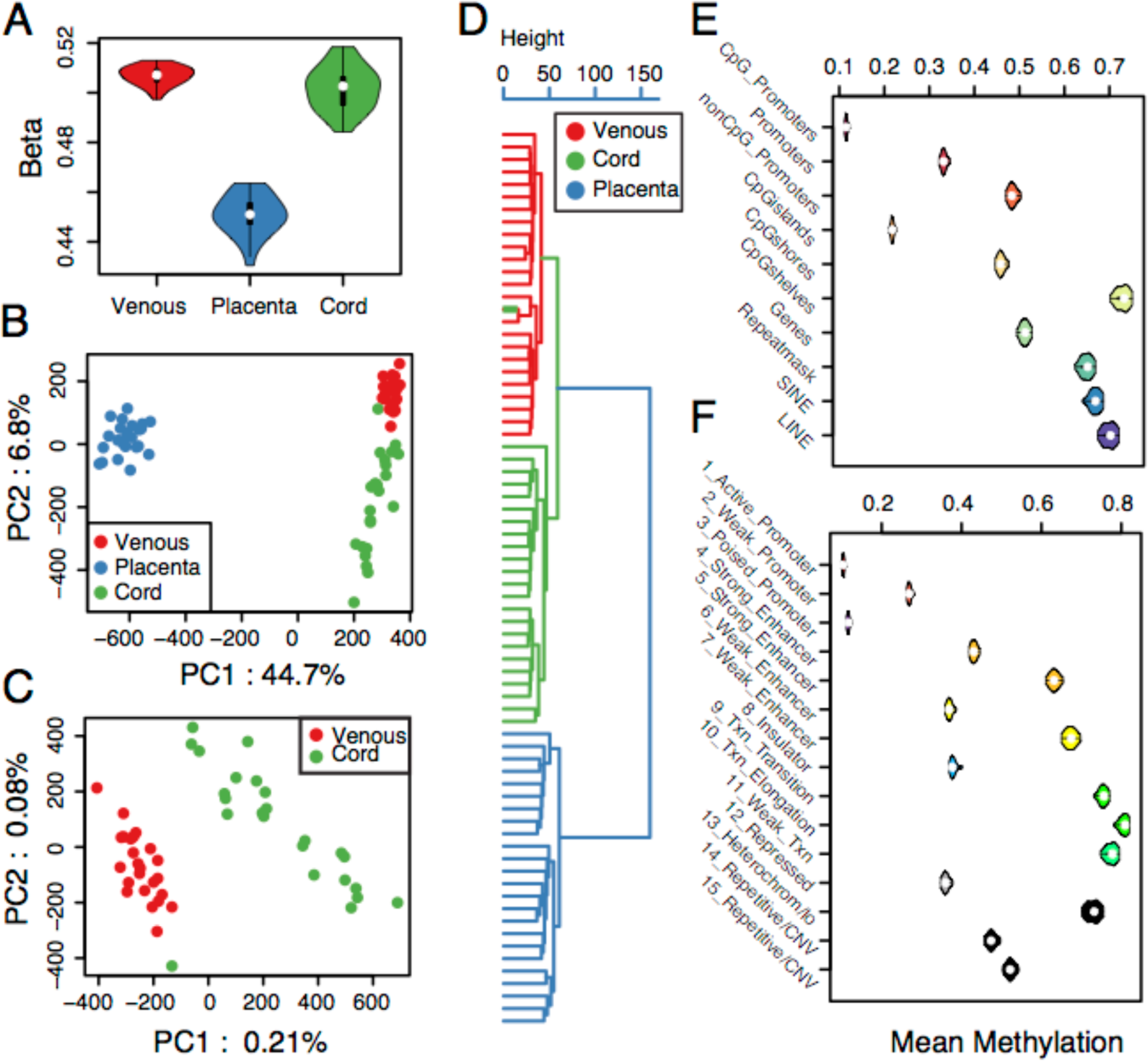
Methylation variation summaries. (A) A violin plot of mean methylation across the entire genome for each individual in each tissue. (B) Dot plot of the first two principle components (PCs) of total methylation variation from all samples. (C) Dot plot of the first two principle components (PCs) of total methylation variation from all blood derived samples. (D) Clustering dendrogram of total methylation variation from all samples. (E-F) Violin plots representing the distribution of mean methylation at all assayed CpG sites partitioned by (E) genomic location and (F) regulatory class in cord blood samples. Box plots for all tissues can be found in supplementary figures (Fig. S1).

Principle component analysis was performed to determine if total genomic methylation variation across all samples could distinguish tissue types (Pai et al. 2011). The first two principle components (PC1 and PC2) both capture methylation variation that is related to tissue type (Fig. 2B). PC1 summarizes 44.7% of the variation and clearly distinguishes placental tissue from the blood tissues, while PC2 captures 6.8% of the variation and largely distinguishes the two blood tissues. Notably, variation among tissues is greater than variation among individuals, consistent with previous work (Pai et al. 2011). The observed broad distribution of cord blood samples led us to evaluate the blood tissue samples in a second independent principle component analysis (Fig. 2C) and again we observed that the first two principle components discriminate tissue type. A single cord blood sample appeared as an outlier that clustered with the venous blood samples (Fig. 2C). To evaluate this finding further, a clustering dendrogram (Fig. 2D) of total genomic methylation variation was constructed. The dendrogram further illustrates the uniqueness of methylation variation among tissue types forming monophyletic clades for each tissue type. The sole exception is the previously observed aberrant cord sample, which pairs with its dyad-pair venous blood sample in the dendrogram, suggesting a potential sample mix-up error; this cord sample was removed from all subsequent analysis.

### Intra-tissue variation is context dependent

Using previously defined functional categories (Ernst and Kellis 2012; Ernst et al. 2011), derived from EnCODE project data (Dunham et al. 2012), each probe was annotated to a putative functional/regulatory class (ChromHMM - see Material and Methods). All probes were also annotated to specific genomic context classes, derived strictly from genome annotations and include promoters, gene bodies, repetitive elements, etc (see Materials and Methods). Both of these annotations were performed so that observed variation could be evaluated by its genomic context and its putative functionality.

Methylation within each tissue varies widely among genome contexts, from ∼10%-70% (Fig. 2E), and is consistent with previous observations (Dunham et al. 2012). Specifically, non-genic regions and repeats are hyper-methylated, gene bodies are in general hemi-methylated while promoters, particularly CpG promoters, are hypomethylated. Further, methylation level increases dramatically with increased distance from CpG islands as indicated by CpG shores and shelf methylation profiles.

Annotation by chromatin state (Faulk and Dolinoy 2011; Ernst and Kellis 2012; Ernst et al. 2011) further illustrates the variability in methylation across classes (Fig. 2F). In general, we expect methylation levels to correlate negatively with chromatin accessibility (Thurman et al. 2012), we expect promoter methylation to have a repressive role on gene expression and we expect genicmethylation to correlate positively with mRNA abundance (Ball et al. 2009). We find that promoter hypomethylation varies by estimated activity, i.e. enhancers are generally hemi-methylated but also vary widely based on class and, as reported previously, on putative gene ontology functions (Barker 1990; Ernst et al. 2011; Gluckman et al. 2010; 2007). Transcribed and functionally inactive regions (Heterochrom/lo) are hypermethylated. Finally, average methylation profiles are consistent across all three tissue types (Fig. S1) suggesting that the defined states will be informative for subsequent analyses.

### Differential methylation across tissues

To identify differential methylation (DM) among tissues (DMT) and quantify the effect of tissue type on methylation variation, we performed a single factor ANOVA and calculated eta-squared (η2) for each CpG site. Eta-squared is defined as the ratio of tissue type sum of squares over total sum of squares. Thus, η2 quantifies the proportion of total methylation variation that can be explained by tissue type. A model including all three tissues (3tissue) revealed that 81.5% of all tested CpG sites were significantly DM at a false discovery rate (FDR) of 5% and yielded a bimodal η2 distribution (Fig. 3A) with a mean η2 of 0.502. In contrast, a model comparing only the two blood tissues (2tissue) found 45.6% of all sites were differentially methylated at an FDR of 5% with a mean η2 of 0.177 (Fig. 3B). Given the bimodality of the 3tissue analysis (Fig. 3A), we looked to determine if the sites at the extremes of the distribution were unique in any way. Indeed, we observe that η2 estimates are influenced by mean methylation with mirrored bimodal distributions for large and small η2 estimates (Fig. 3C). Large η2 sites, those that vary greatly among tissues, are more frequently hypermethylated and small η2 sites more frequently hypomethylated. We return to this observation later.

**Figure 3.**
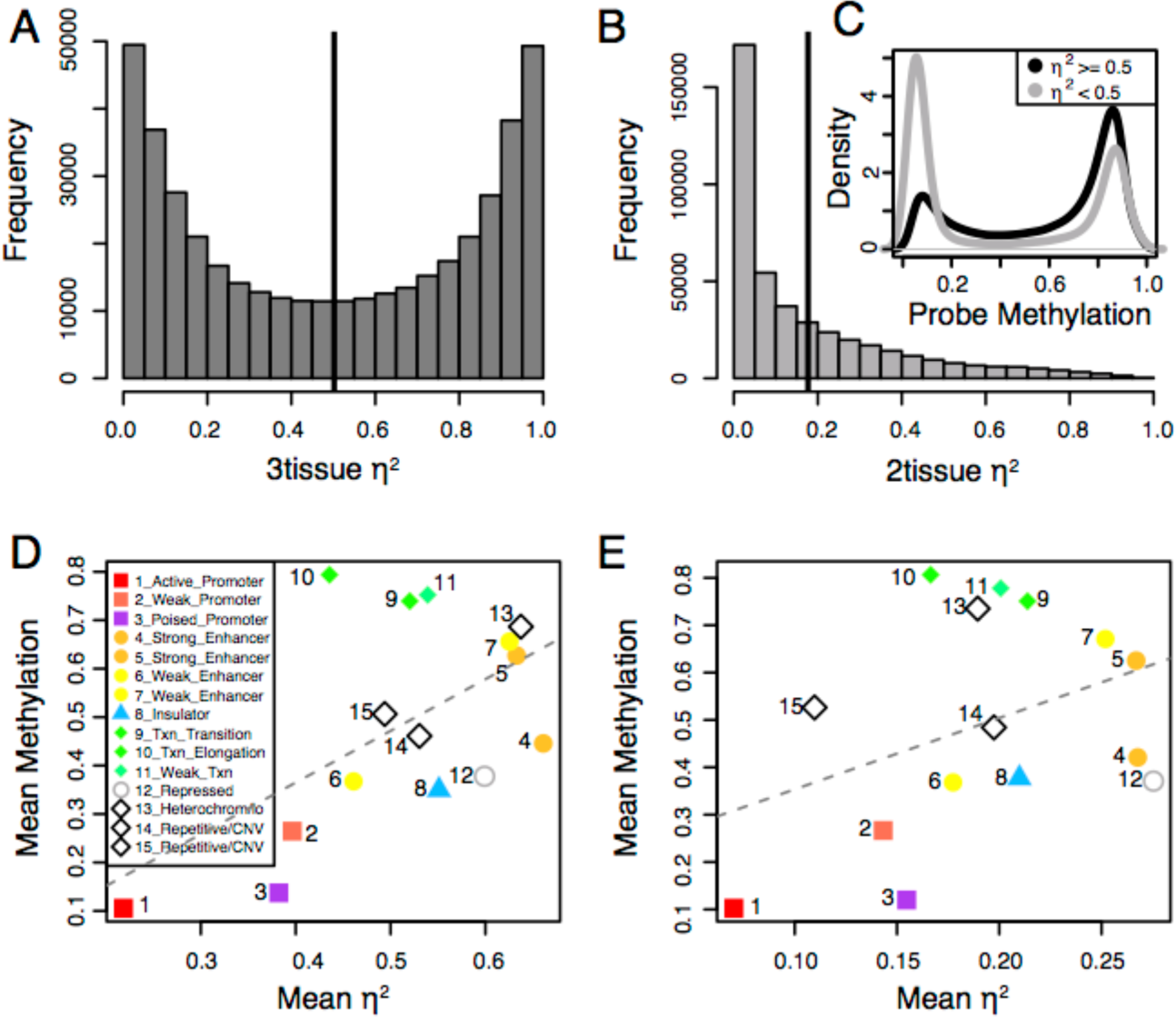
Summary of among tissue Eta-squared (η^2^) estimates. (A-B) The distribution of among tissue η^2^ estimates from (A) three tissues and (B) two tissue analysis. The vertical line in each plot represents the mean of each distribution. (C) Density distribution of methylation levels at CpG sites, averaged across all samples, after partitioning all sites by its 3tissue η^2^ estimates observed in plot (A). (D-E) Mean among tissue η^2^ estimates for each regulatory class and its correlation with mean methylation across all samples. Presented using all three tissues (D) and just the two blood tissues (E).

Next we queried the magnitude of the differences at DM CpGs across tissue types. In the 3tissue model, DM CpGs have, on average, a mean absolute difference of 0.158, between placenta and the two blood tissues. In the 2tissue model, the mean absolute difference is 0.047, revealing that variation in methylation is greater among the placenta and blood tissues than among the blood tissues themselves, consistent with the previously observed difference in total genome mean methylation (Fig. 1A).

### Among tissue methylation is genome context dependent

To determine if η2 estimates have a genome context dependency, we again return to the annotation of 15 previously defined regulatory classes (Ernst et al. 2011; Ernst and Kellis 2012) that include promoters, enhancers, insulators, transcribed, repressed and inactive regions (see Material and Methods). First, we queried if DMT probes were enriched in any specific genomic contexts. We observe an enrichment and depletion in a number of regulatory categories (Table S1). Most striking is the 2.3 and 2.6 fold enrichment observed in class 4 and 5 strong enhancers (p = 2.36e-202 and 1.18e-82, respectively). Insulators are also enriched (1.32-fold, p = 1.81e-16) while poised promoters are depleted (-1.38-fold, p = 1.14e-64). An identical analysis was performed for DMT probes from the 2tissue analysis, which yielded results largely comparable to those performed with the 3tissue data (Table S1).

Next, we averaged η2 estimates for all probes within each functional category and observed large differences across categorical classes. Specifically, promoters, and in particular active promoters, have the least varied methylation profile among tissue types (Fig. 3D). In contrast, strong enhancers have, on average, the most varied methylation profiles across tissue types. This would suggest that methylation variation in regulatory regions outside of promoters might have a larger role in regulating tissue specific gene expression. Again, we repeated the analysis using only the blood tissues (2tissue) and observed congruent results (Fig. 3E).

In an analysis above, we observed that methylation appeared to influence tissue η2 estimates (Fig. 3C). Yet, when binned by functional category, we observe a large difference in mean methylation. As such, we looked to determine what proportion of the variation in methylation could be explained by functional category. To do so, we performed a single factor ANOVA with mean methylation (within tissue) at CpG sites set as the response variable and functional category as the explanatory variable. We estimated that 58.7%, 48.7% and 58.2% of the variation in methylation level could be explained by functional category in cord blood, placenta and venous blood, respectively (F-test p-value = 2.2e-16, for each test). These results suggest that genomic location has a strong influence on methylation level. How then, is among tissue η2 influenced by functional category? To determine this, we again performed a single factor ANOVA with among tissue η2 set as the response variable and functional category set as the explanatory variable. We estimate that 19.8% (F-test p-value = 2.2e-16) of the variation in among tissue η2 can be explained by functional category. These results illustrate the importance of genomic location on the methylation level and tissue specific methylation (and potentially regulatory) profiles.

### Among tissue variation in gene bodies

We repeated the above among tissue η2 analyses, but used mean methylation across gene bodies as the response variable, as opposed to individual CpG sites. Given that gene body methylation is typically positively correlated with gene expression (Ball et al. 2009), we used gene body methylation to more directly determine if differential methylation among tissues is enriched in specific gene ontologies. If validated, this hypothesis would be congruent with gene body methylation having a functional, albeit possibly passive, role (Gutierrez-Arcelus et al. 2013) in tissue or cellular phenotypes. In a 3tissue analysis we observed that 93.8% of all genes are differentially methylated among tissues at an FDR of 5%. To evaluate ontological enrichment, we ranked genes, highest to lowest, by their η2 estimates and tested for enrichment with GOrilla (Eden et al. 2009). In the 3tissue analysis, we observed enrichment (FDR 5%) in 26 biological processes (BPs), 15 molecular functions (MFs) and 8 cellular components (CCs) that consist of genes involved in, but not limited to, the regulation of leukocyte activity, ion transport, response to stimuli, ion channel activity, receptor activity, and located in the plasma membrane part (Table S2). If we invert the genes’ rankings (lowest η2 to highest), thereby asking where is methylation most similar across the three tissues, we then observed enrichment in 32 BPs, 7 MFs and 13 CCs involved in basic cellular functions like gene expression, translation and metabolism and located in, but not limited to, the intracellular and nuclear parts and the ribonucleoprotein complex (Table S3).

In contrast, the 2tissue analysis (cord blood vs venous blood) showed tissue specific differential methylation at 61.7% of genes and enrichment in 163 biological processes, 33 molecular functions and 1 cellular component (transcription factor complex) that can be summarized as involved in embryonic and tissue morphogenesis, tissue development, regulation of gene expression, nucleic acid binding and transcription factor activity, congruent with variation essential for human growth and development (Table S4). These results indicate that gene methylation profiles that are most differentiated among tissues correspond to tissue-specific functions and methylation profiles that are most similar among tissues are related to basic cellular processes.

### Heritability

We estimated the heritability (H) of methylation levels at each CpG site by correlation analysis between maternal and newborn methylation profiles. As both mother and newborns share environmental stimuli, the genetic and environmental influences are confounded. As such, heritability is here defined as the effect of both genotype and environment on methylation. In cord and placental tissue, respectively, we observed that, on average, H is estimated at 0.066 and 0.051 indicating an overall small influence of heritability on methylation variation. However, 4439 (1.03% in cord blood) and 2028 (0.47% in placenta) CpG methylation sites are observed to be heritable at a FDR of 5%, with an H greater than 0.444. Furthermore, even though only a small percentage of sites may be defined as heritable, we observe a 30.5% overlap in heritable sites between cord blood and placental tissues (Fig. 4A). To identify any potential impact of heritable methylation levels on phenotypes, we queried their genomic context (union of all H CpGs = 4954) and found that heritable methylation profiles were observed in all regulatory categories. However, a paucity of heritable methylation in all three of the promoter categories and an enrichment of heritable methylation in insulators, strong enhancers as well as in weakly transcribed, inactive and repetitive categories was observed (Table 2). Congruent with our analyses of DM among tissues, these observations highlight the uniqueness of methylation variation outside of promoter regions.

**Figure 4.**
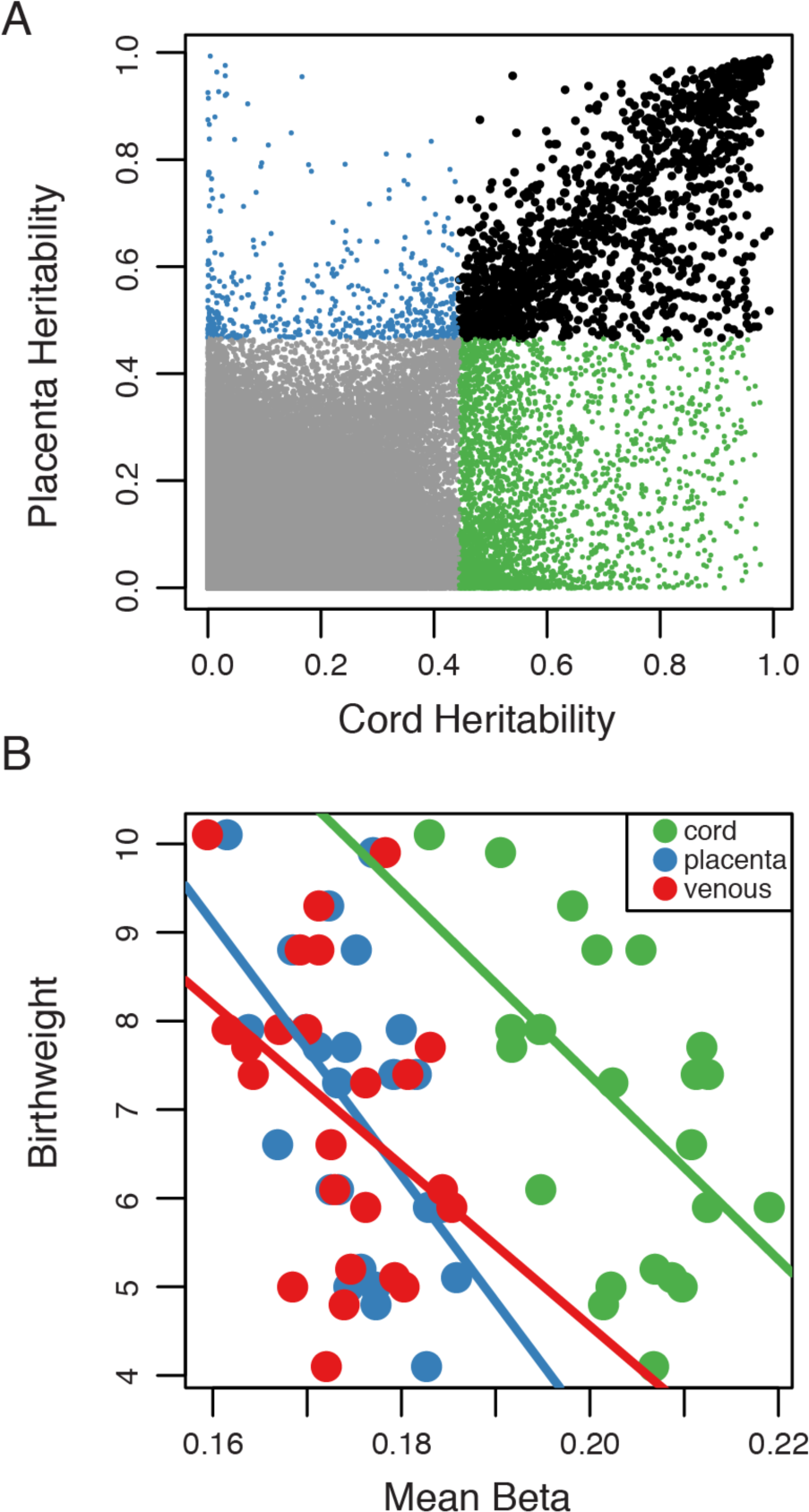
Methylation heritability and *MTHFR*-birthweight correlations. (A) Dot plot of the correlation in heritability estimates between cord-venous and placenta-venous analysis. Points in green represent all sites observed to be significantly heritable in cord blood. Points in blue represent all sites observed to be significantly heritable in placenta tissue. Points in black represent sites observed to be significantly heritable in both placenta tissue and cord blood. Significance called at an FDR of 5%. (B) Dot plot of the correlation between mean *MTHFR* gene body methylation and newborn birthweight within each tissue.

**Table 2.** Regulatory Enrichment for Heritable Probes

| Functional Category | Enrichment | padj |
| --- | --- | --- |
| 1_Active_Promoter | -2.44 | 2.73E-84 |
| 2_Weak_Promoter | -1.40 | 2.49E-06 |
| 3_Poised_Promoter | -2.48 | 8.13E-20 |
| 12_Repressed | -1.18 | 1.01E-02 |
| 4_Strong_Enhancer | 1.26 | 4.22E-02 |
| 8_Insulator | 1.58 | 2.64E-05 |
| 11_Weak_Txn | 1.24 | 3.32E-05 |
| 13_Heterochrom/lo | 1.55 | 2.79E-49 |
| 14_Repetitive/CNV | 2.71 | 1.63E-04 |

Functional categories, enrichment estimates and Bonferroni corrected p-values (padj) for heritable probes.

### Global methylation and stress

We next tested for an association between maternal stress and average genome-wide methylation level, or simply the average methylation level across an individual-x-tissue’s genome. Strikingly, genome wide mean methylome profiles in maternal venous blood are correlated with maternal stress (Fig. S2A, r = −0.608, p = 0.002), a result not observed in either of the newborn tissues. Furthermore, mean methylome variation in maternal venous blood is also correlated with rape occurrence (r = −0.618, p = 0.001), newborn birthweight (r = 0.601, p = 0.002), maternal weight (r = .426, p = 0.043), maternal age (r = 0.494, p = 0.017) and household education (r = 0.419, p = 0.041) (Fig. S2B-F). As observed previously in the correlation of stress and other factors on newborn birthweight, the observed correlations with mean methylation in venous blood are also confounded by multiple explanatory variables. However, these observations may be the product of cell type heterogeneity in maternal venous blood as opposed to actual alterations in methylation across the genome (Ožura et al. 2012; Dhabhar et al. 2012; Gotovac et al. 2010; Vidović et al. 2007; Boscarino and Chang 1999; Amati et al. 2010; Atanackovic et al. 2006).

### Local methylation and stress

We estimated the effect of stress on methylation at three genomic levels - (1) methylation at each CpG site and an average of CpG methylation at each (2) promoter and each (3) gene body. A multi-factor mixed model ANOVA was performed and a η^2^ was calculated for each factor (see Materials and Methods). The model includes a fixed tissue effect and a random mean methylome estimate effect to account for possible cell type heterogeneities. After Benjamini Hochburg correction of the p-value distributions, we observed 212, 13, and 37 CpGs, promoters and genes, respectively, significantly correlated with maternal stress at a FDR of 5% (315, 24, 71, respectively, at a FDR of 10%). The average proportion of explained variation (η^2^) for stress at significant CpGs, promoters and genes was estimated at 19.7% (max = 66.1%), 14.6% (max = 29.4%) and 14.3% (max = 39.0%), respectively.

Interestingly, 43.4% of stress related sites, i.e. those significant at an FDR of 5%, were also observed to be heritable at an FDR of 5%. This ratio increases to 75.9% if we consider stress related probes at an FDR of 20%. In other words, CpG sites that are correlated with maternal stress are also largely identified as heritable. This observation is consistent with a hypothesis that stress-mediated changes in methylation may be heritable.

To identify pathways that may be influenced by stress, we ranked all promoters and genes by their η^2^ estimates and tested for enrichment in gene ontologies using GOrilla. From promoters, we observed enrichment in four cellular component pathways involved in intracellular and organelle parts and with genes, we observed enrichment in similar cellular component pathways as well as in 21 molecular function pathways involved in protein ubiquitination, cellular metabolism and mitotic cell cycle (Table S5 and Fig. S3).

### Stress-related methylation variation is correlated with birthweight

To determine if stress-related methylation variation may have an effect on an organismal phenotype, specifically birthweight, we performed a single factor ANOVA analysis using methylation at the 212 stress-related CpG sites (5% FDR) as the explanatory variable and birthweight as the response variable. After Bonferroni correction, 13 CpG sites (Table 3) exhibited a significant effect and are associated with 10 genes involved with processes that include E3 ubiquitin ligase, calcineurin signaling and multiple developmental processes.

**Table 3.**
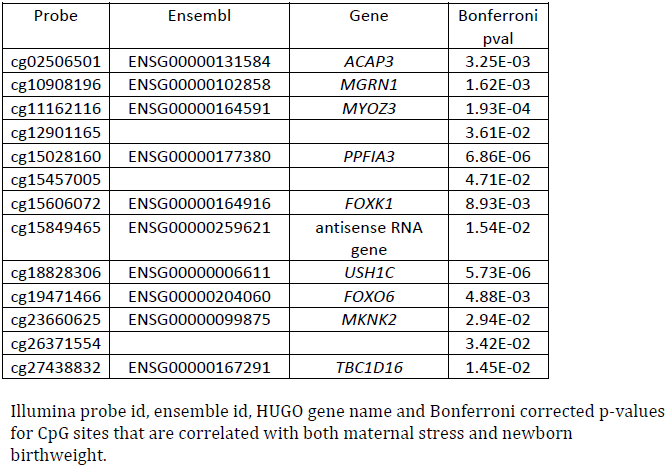
CpG Sites Correlated With Both Stress and Birthweight

If we consider stress associated promoters and genes, rather than CpG sites, one promoter (*SAC3D1*) and two gene bodies (*SLC29A1* and *MYOZ3*, also seen in the CpG site analysis) also exhibited a significant correlation with birthweight after Bonferroni correction.

### Methylation and birthweight

In a separate analysis, we performed a three factor mixed model ANOVA identical to that performed with methylation and stress, with the exception that we replaced stress with birthweight, i.e. we tested for an association between methylation and birthweight. The model was fit against methylation at all CpG site, promoters and gene bodies. We observed six CpG sites, six promoter regions and six genes with a significant birthweight effect at a FDR of 5% (Table S6). Among the associated genes are *PPF1A3*, *USH1C* and *MYOZ3*, each observed in the stress-related analyses above.

One gene that was not observed in previous analyses is *MTHFR* (methelenetetrahydrofolate reductase), a gene necessary for proper folate metabolism and previously associated with global DNA methylation (Friso 2002 PNAS), neural tube defects (van der Put 1995), depression (Bjelland, 2003), bi-polar disorder (Gilbody, 2007) and schizophrenia (Zhang 2013). In this study, we found that an increase in *MTHFR* gene body methylation is associated with a decrease in birthweight. When testing each tissue independently, we observed a significant correlation between *MTHFR* methylation in placenta (r = −0.52, p = 0.009) and cord blood tissues (r = −0.55, p = 0.006) with birthweight, but not in venous blood (r = - 0.38, p = 0.067) (Fig. 4B).

## Discussion

We used a unique collection of human tissues (maternal venous blood, newborn cord blood, and placenta) derived from new mothers who have experienced different levels of extreme physical and psycho-social stress to evaluate the role that stress, heritability, genomic context and tissue type play in shaping genome-wide methylation variation and outcome phenotypes like newborn birthweight.

Genome wide methylation profiles vary greatly among assayed tissues, separating on the first two principle components and forming unique monophyletic clades in an unsupervised hierarchical clustering analysis. Tissue specific methylation was observed at the majority of assayed sites among the three tissues (81.5%), suggesting that methylation variation in all genomic contexts plays an important role in tissue-specific phenotypes. However, the putative magnitude of that effect varies by genome context. Specifically, our results support a hypothesis that methylation outside of promoter regions has a larger average effect on tissue and cellular phenotypes than methylation variation within promoter regions. This result does not diminish the importance of promoters since extreme eta-squared values are observed in all contexts (Fig. S4). Rather, regions other than promoters, such as enhancers and insulators, have more dynamic methylation profiles and may resultantly have a larger effect on tissue/cellular phenotypes. As such, studies that limit their investigation to promoter regions may be missing more influential methylation variation elsewhere.

A recurrent question in studies of methylation is the heritability, or specifically the additive genetic component of heritability, of methylation. This is often assessed by identifying differential methylation among offspring derived from parental groups who were exposed to different treatments (Sapolsky et al. 2000; Carone et al. 2010; Herman et al. 2003; Kalantaridou et al. 2004) but also through the direct correlation of parent and offspring methylation profiles (Fraser et al. 2012). In our system, the genetic and environmental components of heritability are confounded as we assume that the in utero tissues are influenced by both the mother’s environment and the maternal response to said environment, in addition to maternal and paternal genetic contributions to methylation. However, in comparison to other studies, our heritability estimates were comparably low, with an overall mean H of ∼0.06 and significant heritability (H > 0.45) observed at roughly 1% of sites. Nonetheless, where heritable methylation was observed, it was enriched in insulators and enhancers, among other regions, and depleted in promoters. Again, this demonstrates a putatively larger role in gene regulation for non-promoter methylation variation.

Finally, we used our system to determine if methylation variation may mediate the effect of stress on large organismal phenotypes, specifically in this case newborn birthweight. First, we observed that stress, maternal age, education, and birthweight were inter-correlated. We propose that in this sample, i.e. women in the DRC, age and household education may influence, and possibly reduce, maternal stress. For example, increased age and education may enable a larger social network and social support that would mitigate the effects of stress, a proposal consistent with our finding that increased age and education is correlated with decreased stress. Thus, we hypothesize that it is the physical properties of war stress acting through the hypothalamic-pituitary-adrenal axis that influence methylation variation and newborn birthweight.

Following this hypothesis, we tested for an association between stress and methylation and then methylation and newborn birthweight. We first observed that our model of maternal stress was correlated with mean global methylome variation in venous blood and in turn mean venous blood methylome variation was correlated with newborn birthweight (r = −0.601, p = 0.002). Previously, we found a correlation between maternal stress, newborn methylation at the glucocorticoid receptor *NR3C1* and newborn birthweight (Mulligan et al. 2012). Given these results, we hypothesize that maternal stress is correlated with genome-wide changes in methylation only in the mother (venous blood) because the mother most directly experiences the stressors whereas the effect of maternal stress on newborn methylation is reflected only in specific genes associated with the response phenotype (birthweight), such as *NR3C1*.

An important noteworthy limitation of our study, as with all studies profiling methylation variation in a tissue, is cell type heterogeneity. Specific to venous blood, previous studies have observed stress-induced changes in cell type composition in whole blood (Ožura et al. 2012; Dhabhar et al. 2012; Gotovac et al. 2010; Vidović et al. 2007; Boscarino and Chang 1999; Amati et al. 2010; Atanackovic et al. 2006). As such, observed alterations in methylation could alternatively be explained by alterations in cell type compositions. Because of this possibility, all fitted models of methylation presented here included mean methylation, along with tissue type, as co-factors.

As a product of the observations discussed above, we developed a model of methylation variation to estimate the influence of tissue type, mean methylome level, and stress on each CpG, each promoter and each gene body. Estimates of explained variation were summarized with the η^2^ statistic. Overall, a small fraction of sites, promoters and genes were significantly correlated with stress. However, among the stress-correlated genes, there was an observed enrichment in protein ubiquitination and metabolism pathways that are influential in numerous cellular phenotypes, including histone modification (Shilatifard 2006), inflammatory response (Wang and Maldonado 2006), human disease (Kessler 2013) and cell growth and division (Benanti 2012). In addition, genetic variation in ubiquitin-related genes has been previously associated with growth in yeast (Roscoe et al. 2013), cancer cells (Yang et al. 2013), chickens (Sheng et al. 2013) and cattle (Wang et al. 2013). These observations suggest that stress-associated methylation variation may mediate fetal growth, and thus birthweight, as well as influence later physical and mental health in the offspring and current maternal health. Interestingly, these observations are consistent with other results and hypotheses on the influence of periconceptual environment and glucocorticoid exposure on fetal metabolism and growth (Elizabeth C Cottrell 2009; Hussain 2012; Bolten et al. 2011; Seckl 2006; Seckl and Holmes 2007). Additionally, a recent paper found no significant differences in global methylation levels between cord blood and three year old children, suggesting that factors that influence DNA methylation may confer long-term effects (Herbstman et al. 2013). Thus, our results support the hypothesis that environmentally-induced methylation modifications may mediate maternal disease as well as fetal phenotype and future disease risk. Finally, these observations illustrate the importance of decreasing violence against women and advocating for women’s equality worldwide. As implicated here and by the World Health Organization (WHO et al. 2013), violence against women does not just influence their physical and emotional health, but potentially that of future generations as well.

## Materials and Methods

### Samples

Whole blood samples were collected from 24 women who delivered their babies at HEAL Africa hospital in Goma, eastern Democratic Republic of Congo in July-August, 2010. Umbilical cord blood and placenta samples were collected from the same 24 women within several hours of delivery (Mulligan et al. 2012). Genomic DNA was isolated on site using Qiagen QIAamp DNA Midi Kits (Qiagen, Valencia, CA) according to the manufacturer’s instructions. Samples were collected with informed consent under Western IRB approval, Olympia, WA (WIRB Project # 20100993), the University of Goma, and from an ethical review committee at HEAL Africa hospital.

### Ethnographic and trauma interviews

Trauma surveys and ethnographic interviews were conducted with each of the 24 participants within 24 hours of giving birth to collect data on past and culturally relevant physical and psycho-social (Table 1). Trauma surveys were adapted and expanded for cultural relevance from the Peritraumatic Distress Inventory (Brunet et al. 2001). Ethnographic interviews (Spradley 1979) were conducted with each participant to cover reproductive history, past traumatic exposures, and general health history. Anthropomorphic data were collected on mothers (height, weight) and newborns (birthweight, birth length, gestational age).

### DNA Methylation Data

Isolated and purified genomic DNA was shipped to the University of Miami Hussman Institute for Human Genomics in Miami, Florida where 500 ng of each sample was bisulfite converted and processed on Illumina HumanMethylation450 BeadChips following the manufacturer’s specifications. Prior to shipment, all samples were randomized and given a unique tissue and dyad blind identifier, simply numbered 1 through 72, to insure that all samples were processed in a non-structured manner. Raw data were processed through Illumina’s GenomeStudio V2011.1 Methylation Module v1.9.0 for cross array normalization and output of average Beta estimates and detection p-values. Any probe with a detection p-value greater than 0.01 in any one sample was eliminated from further analysis along with all probes located on the sex chromosomes. We observed that approximately 5% of the methylation data failed a Shapiro-Wilk normality test, however no transformation (log2, logit, N(0,1)) of the data improved this proportion. As such no transformation of Beta values was performed.

### Annotation

A new annotation was performed for each probe mapping their genomic location to genes, promoters, CpG islands, CpG shores, CpG shelf, SINEs, LINEs and additionally to chromatin defined regulatory states (Ernst et al. 2011; Ernst and Kellis 2012), which includes promoters, enhancers, insulators, transcribed, repressed and inactive regions. Ensemble v69 gene, CpG island and Repeatmasker data was downloaded from UCSC v37 on Dec 12, 2012. Promoters were defined as the first 2000 base pairs 5’ of the transcription start site. CpG shores were defined as the first 2000 base pairs both 5’ and 3’ of a CpG island and CpG shelves were defined as the next 2000 base pairs 5’ and 3’ of CpG shores. In cases of neighboring CpG island-shore-shelf overlaps, a probe would be annotated into the highest annotation class only. Chromatin regulatory states were defined using B-lymphoblastoid (GM12878) data from Ernst et al (Ernst et al. 2011). Fifteen states are defined within six broader states, namely: promoter, enhancer, insulator, transcribed, repressed and inactive regions. We note that while data from a B-lymphoblastoid cell line may currently provide the best model for the whole blood cord and venous tissues, it may not fairly represent chromatin states in placental tissue. However, with no other suitable cell type data currently available, and in an effort to compare and contrast identical regions, we proceeded cautiously.

### Statistical Analysis

All statistical analyses were carried out in R using in-house scripts. A commonly used statistic is η^2^ (Eta-squared), which quantifies the proportion of variation explained by a modeled factor of interest. It is defined as the sum of squares for your factor of interest over the total sum of squares.

To estimate the effect of stress on CpGs, we performed a mixed model ANOVA including mean methylation and tissue as co-factors in the model. The model is defined as: methylation = a + methylome + tissue + stress + stress*tissue + error, where “a” is the fixed average methylation level for the modeled CpG site, “methylome” is the random genome wide methylation average for a sample, “tissue” is the fixed tissue effect, “stress” is the random effect due to environmentally induced war stressors, “stress*tissue” is the random effects due to interaction or altered responses to stress among tissues and “error” accounts for all unexplained or residual variation.

Identical analyses were performed on promoters and gene bodies as described above for CpG sites. However, methylation at each promoter or gene body was summarized as the mean methylation for all CpG sites within the boundaries of the analyzed promoter or gene.

A beta regression analysis was also performed on a subset of the data using the betareg function in R. Significance was determined by fitting two models, a null model with tissue type as an explanatory variable and mean genomic methylation as an offset, and a second model was then fit with stress as an additional explanatory variable. Significance was determined by log-likelihood test. A small increase in the number of significant stress-related methylation profiles was observed and ∼95% of our significant profiles from the linear regression were replicated. Given the additional information we gain by calculating the eta-squared statistic (η^2^) from the more simplistic linear model, we continued to use the linear model over the beta regression.

## Data Access

All Illumina Methylation450 K array data has been deposited in Gene Expression Omnibus (GEO) under accession number GSE54399.

## Acknowledgments

We thank the women of the Democratic Republic of Congo for their participation in this study. Funding was provided by NSF grant BCS 1231264 and grants from the University of Florida (UF) Clinical and Translational Science Institute, UF College of Liberal Arts and Science, and a UF Research Opportunity Seed Fund award.

